# Rates of global cellular translation and transcription during cell growth and the cell cycle in fission yeast

**DOI:** 10.1101/2022.10.17.512486

**Authors:** Clovis Basier, Paul Nurse

## Abstract

Proliferating eukaryotic cells grow and undergo cycles of cell division. Growth is continuous whilst the cell cycle consists of discrete events. How the production of biomass is controlled as cells increase in size and proceed through the cell cycle is important for understanding the regulation of global cellular growth. This has been studied for decades but has not yielded consistent results. Previous studies investigating how cell size, the amount of DNA, and cell cycle events affect the global cellular production of proteins and RNA molecules have led to highly conflicting results, probably due to perturbations induced by the synchronisation methods used. To avoid these perturbations, we have developed a system to assay unperturbed exponentially growing populations of fission yeast cells. We generated thousands of single-cell measurements of cell size, of cell cycle stage, and of the levels of global cellular translation and transcription. This has allowed us to determine how cellular changes arising from progression through the cell cycle and cells growing in size affect global cellular translation and transcription. We show that translation scales with size, and additionally increases at late S-phase/early G2, then increases early in mitosis and decreases later in mitosis, suggesting that cell cycle controls are operative over global cellular translation. Transcription increases with both size and the amount of DNA, suggesting that the level of transcription of a cell may be the result of a dynamic equilibrium between the number of RNA polymerases associating and disassociating from DNA.

## Introduction

Proliferating steady state eukaryotic cells undergo two fundamental processes: they increase in biomass and they undergo cycles of cell division. Biomass increase is a continuous process whilst the cell cycle consists of an orderly transition through a series of specific discrete events. How these continuous and punctuated processes impact upon the accumulation of proteins and RNA, the major drivers of biomass increase, is important for understanding how overall cellular growth is regulated [1]. Proteins make up 35-60 % and RNA 4-12 % of the dry mass of cells [2] and their production consumes more than half of the ATP of a cell [3]. Previous studies of the patterns of protein and RNA through the cell cycle have led to highly conflicting results. In this paper we have addressed this problem using unperturbed steady state growing fission yeast cells.

Knowing the pattern of protein and RNA accumulation during the growth of cells though the cell cycle is an example of the general problem of scaling, a power-law relationship between two variables. The accurate scaling of protein and RNA synthesis with cell size maintains their concentration at a constant level, and there is evidence that loss of proper scaling of biosynthesis leads to cellular dysfunction and may be a causal driver for aging and senescence [4–7]. Growth is continuous whilst cell cycle events are temporally discrete changes within cells. In particular, the amount of DNA, the template for RNA production, doubles once during S-phase early on in the cycle, and mitosis and cell division at the end of the cell cycle involve major cellular reconstruction. These can affect global cellular translation, the rate of synthesis of all proteins, and global cellular transcription, the rate of synthesis of all RNA molecules.

Global cellular translation has been investigated in numerous systems with varying results. Studies using incorporation of exogenous amino acids to measure global translation in populations of synchronised yeasts have yielded conficting results, either that global cellular translation undergoes significant changes during the cell cycle [8] or remains constant [9–11]. In mammalian cell cultures, studies monitoring global cellular translation through synchronised cell cycles were also contradicting, with some finding no changes [12] whilst others found increases and/or decreases of varying magnitudes during mitosis [13–18]. Asynchronous cultures of yeasts and mammalian cells have not detected major cell-cycle related changes suggesting that previous discrepancies may have been due to synchronisation methods [11]. Previous studies of global cellular transcription through the cell cycle have relied on population measurements of the incorporation into RNA of pulse-labelled nucleobases or nucleosides in synchronous cultures. In the two yeasts, *Schizosaccharomyces pombe* [19–23], and *Saccharomyces cerevisiae* [9,24–26], these studies have yielded variable results. Some studies found that RNA synthesis increased at a discrete stage of the cell cycle, either at DNA replication [19,20,25] or later [21–23], whilst others found a constant increase throughout the cell cycle [9,24,26]. It is likely that these discrepancies arise from the different protocols used to generate synchronous populations [23]. Work in unperturbed mammalian cell lines suggests that global cellular transcription increases from G1 to G2 [27].

In this work, we characterise the scaling of global cellular translation and global cellular transcription through the growth of cells and progression throughout the cell cycle using single-cell approaches in steady-state exponentially growing fission yeast cells to avoid problems induced by cell-cycle synchronisation.

## Results

### Single-cell assays to measure global cellular translation and transcription in asynchronous steady state exponentially growing cultures

To measure rates of global cellular translation and transcription through the cell cycle of fission yeast cells, we developed assays to quantify these rates whilst measuring cell size and identifying cell cycle stages in thousands of single cells in exponentially growing cultures. This is possible because fission yeast cells are rods that grow by tip elongation, so cell length is an indicator of cell cycle position [28].

To quantify global cellular translation, we incubated cells with a methionine analogue, L-homopropargylglycine (HPG), and measured its incorporation into proteins using Click Chemistry. Wild type cells were incubated for 5 minutes with HPG, and were stained with an Alexa Fluor azide (Figure 1A, B). The increase in HPG labelling had almost no lag and was essentially linear for a 5-minute period after HPG addition (Figure 1C), indicating that a 5-minute pulse could be used to estimate the rate of HPG incorporation. The pulse signal was five times the background signal. Digestion of protein molecules using Proteinase K removed the fluorescent signal (Figure S1A), and inhibiting translation using cycloheximide inhibited HPG incorporation (Figure S1B). Thus, a 5-minute HPG incubation and labelling can be used as a measure of global cellular translation. There is some HPG signal in the nucleus (Figure 1B), which is possibly the result of nuclear translation [11,29,30] and/or rapid translocation into the nucleus of peptides synthesised in the cytoplasm.

**Figure 1.**
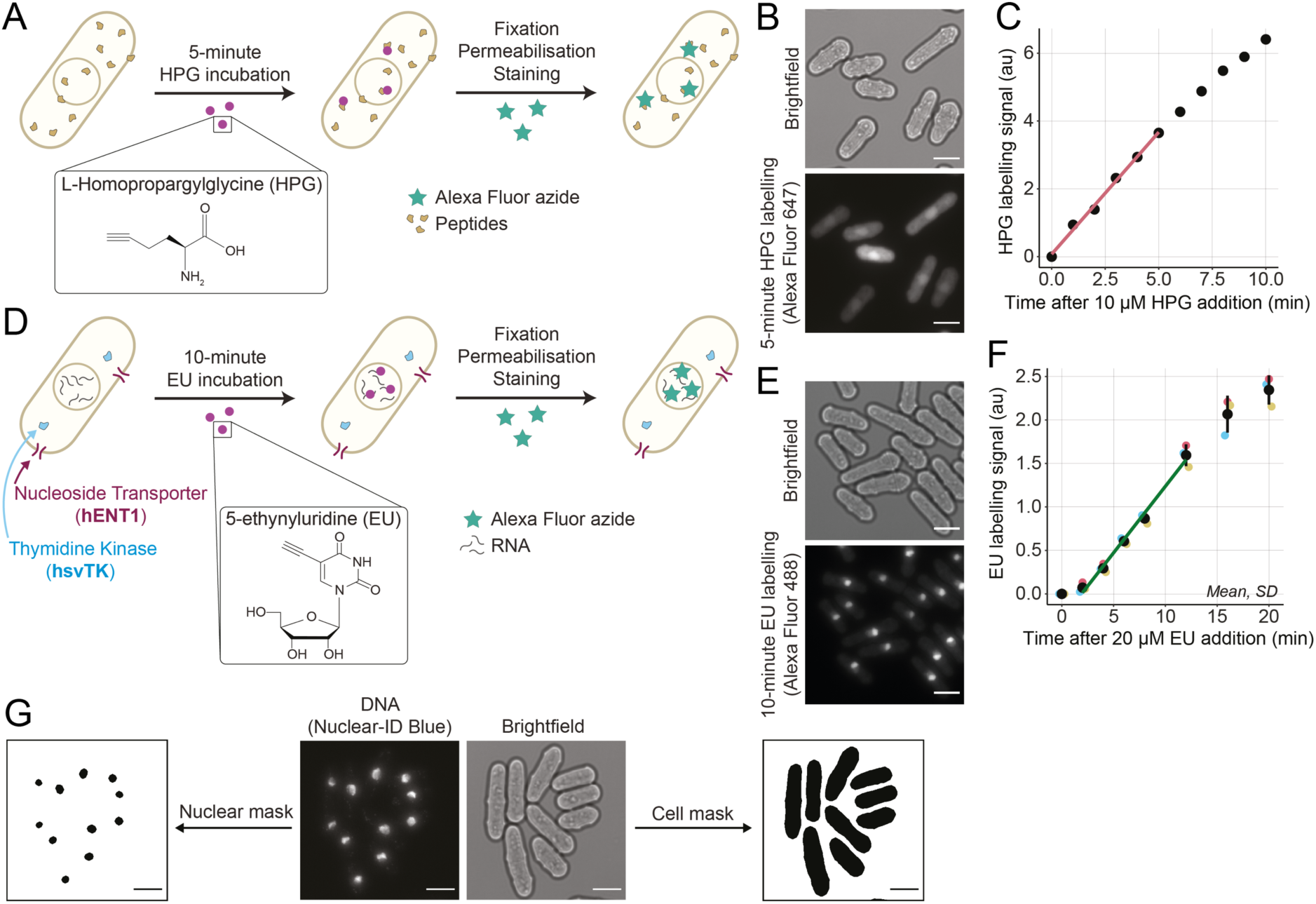
Single-cell assays to measure global cellular translation and transcription in steady state growing asynchronous cultures. (**A**) Overview of the global cellular translation assay. Wild type cells are incubated with HPG for 5 minutes, then fixed, permeabilised, and an Alexa Fluor azide fluorophore is covalently attached to HPG molecules using Click Chemistry. (**B**) Example images of brightfield and fluorescently labelled HPG (Alexa Fluor 647) of wild type cells assayed for global cellular translation. Scale bars represent 5 µm. (**C**) Change in HPG labelling signal with different durations of HPG incubation, measured by flow cytometry. Population medians of at least 200,000 cells are shown. The red line is the ordinary least square (OLS) linear regression fitted on the medians between 0 and 5 minutes. (**D**) Overview of the global cellular transcription assay. Cells expressing *hENT1* and *hsvTK* are incubated with EU for 10 minutes, then fixed, permeabilised, and an Alexa Fluor azide fluorophore is covalently attached to EU molecules using Click Chemistry. (**E**) Example images of brightfield and fluorescently labelled EU (Alexa Fluor 488) of *hENT1* and *hsvTK* cells assayed for global cellular transcription. Scale bars represent 5 µm. (**F**) Change in EU labelling signal with different lengths of EU incubations, measured by flow cytometry. The mean and standard deviation (SD) of the population medians of at least 200,000 cells in experimental triplicates are shown in black. The dark green line is the OLS linear regression fitted on the mean data between 2 and 12 minutes. (**G**) Example images of brightfield used to generate cell masks, and DNA (Nuclear-ID Blue) used to generate nuclear masks. Scale bars represent 5 µm.

To quantify global cellular transcription, we incubated cells with the uridine analogue, 5-ethynyluridine (EU) and measured its incorporation into all major RNA species [31] using Click Chemistry to fluorescently label EU molecules. EU was added to a culture of exponentially growing cells expressing the human Equilibriative Nucleoside Transporter 1 (hENT1) and the herpes simplex virus Thymidine Kinase (hsvTK), necessary for the uptake and phosphorylation of EU (Figure S1C). After 10 minutes, cells were fixed, permeabilised, and EU molecules were fluorescently labelled with an Alexa Fluor azide (Figure 1D, E). The EU labelling signal was linear from 2 min to 12 min (Figure 1F) indicating that a 10-minute incubation could be used to estimate the rate of EU incorporation into RNA. Linearity is also not much influenced by a longer 20-minute incubation. However, in contrast to the translation assay, the transcription pulse signal was less strong and was only twice the background signal so may be a less accurate estimate of the rate of transcription. Digestion of RNA molecules using RNAse A removed the EU labelling signal (Figure S1D), and inhibition of RNA polymerases using 1,10-phenanthronline inhibited EU incorporation (Figure S1E). Thus, a 10/20-minute EU incubation and labelling can be used as a measure of global cellular transcription. Although the precise fractions of the different types of RNA in global transcription have not been fully characterised, recent work indicated that only half of the newly synthesised RNA consists of ribosomal RNA molecules, suggesting that a significant portion of transcription is dedicated to the production of messenger and other RNA molecules [27].

To obtain single-cell measurements of cell size and global translation or transcription, we used brightfield and fluorescence microscopy in combination with automated segmentation tools [32]. A fluorescent DNA dye (Nuclear-ID Blue) was used to identify and remove binucleated cells allowing analysis of a cell population in which all cells are composed of a single nucleus (Figure 1G).

### Global cellular translation and transcription change with cell length

We used these two assays to investigate how global cellular translation and transcription are affected by cell size, and by progression through the cell cycle within an asynchronous population (Figure 2A, D). We used cell length as a measurement of size, as cell length is correlated with cell volume because *S. pombe* cells are cylinders that grow by tip extension [28]. Binucleated and septated cells were excluded from the analysis to eliminate the effects of mitosis and cell division, as well as S-phase which occurs during septation in wild type cells [33]. The median global translation increased smoothly with cell length (Figure 2B) and global translation per cell length was found to be constant with size (Figure 2C, S2A, S2B). Thus, in the wild type cells, global translation scales linearly with cell size as mononucleated cells proceed through the cell cycle.

**Figure 2.**
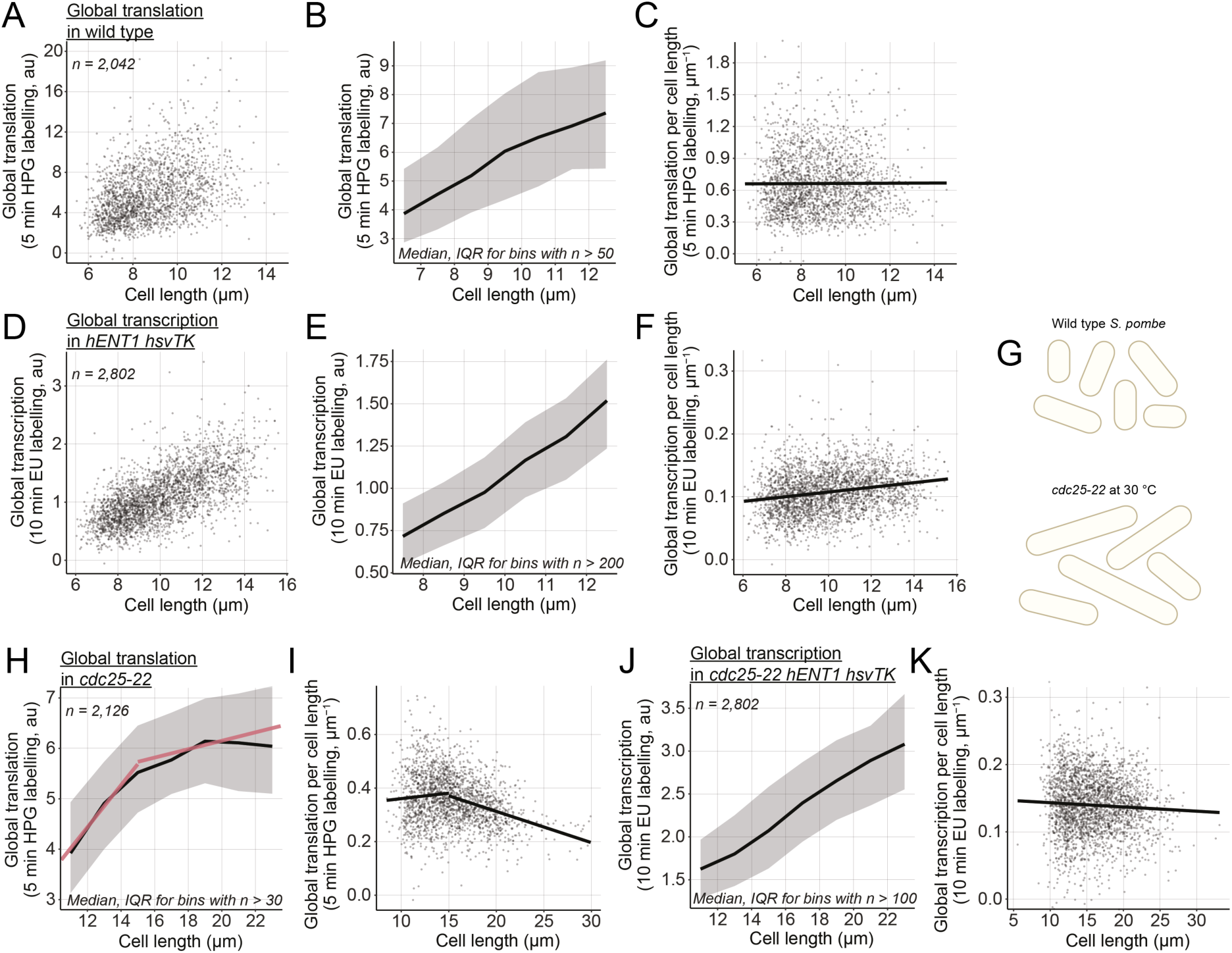
Global cellular translation and transcription with cell length in wild type cells. (**A**) Global cellular translation of wild type single cells. (**B**) Medians of global translation (solid black line) and IQR (shaded area) of cells shown in (A) grouped in length bins of 1 µm. Bins containing more than 50 cells are shown. (**C**) Global cellular translation of cells shown in (A) divided by their cell length. The solid black line represents the OLS linear regression fitted on the data. (**D**) Global cellular transcription of single cells expressing *hENT1* and *hsvTK*. (**E**) Medians of global transcription (solid black line) and interquartile ranges (IQR, shaded area) of cells shown in (D) grouped in length bins of 1 µm. Bins containing more than 200 cells are shown. (**F**) Global cellular transcription of cells shown in (D) divided by their cell length. The solid black line represents the OLS linear regression fitted on the data. (**G**) Schematic representation of cell length in asynchronous populations of the wild type and the *cdc25-22* mutant grown at the semi-permissive temperature (30 °C). (**H**) Medians of global translation (solid black line) and IQR (shaded area) of *cdc25-22* cells grown at 30 °C and grouped in length bins of 2 µm. Bins containing more than 30 cells are shown. The solid red lines represent OLS linear regressions fitted on the single-cell data for cells shorter and longer than 15 µm. (**I**) Global cellular translation of cells shown in (H) divided by their cell length. The solid black line represents the OLS linear regression fitted on the data for cell shorter and longer than 15 µm. (**J**) Medians of global cellular transcription (solid black line) and IQR (shaded area) of *cdc25-22 hENT1 hsvTK* cells grown at 30 °C and grouped in length bins of 2 µm. Bins containing more than 100 cells are shown. (**K**) Global cellular transcription of cells shown in (J) divided by their cell length. The solid black line represents the OLS linear regression fitted on the data.

Likewise, median global transcription increased smoothly with cell length in the *hENT1 hsvTK* population (Figure 2E), although global transcription per cell length increased somewhat as cell length increased during the cell cycle (Figure 2C, S2A, S2B). Because the cells are growing in steady state it would be expected that the rate per unit cell length at the end of the cell cycle would be the same as at the beginning of the cell cycle. We speculate that the increase we observed may be a consequence of the low signal to noise ratio of around 2:1 leading to a technical defect in the background estimate. However, we can conclude that the rise in transcription as cells grow does not exhibit any discontinuities.

To investigate the effect of sizes beyond those of wild type cells, we used the temperature sensitive *cdc25-22* allele. When grown at a semi-permissive temperature, *cdc25-22* cells divide at longer lengths than wild type cells whilst maintaining the same growth rate and not displaying any cell cycle defects (Figure 2G) [34, 35]. We found that global translation increased with cell length only in cells up to a length of 15 µm, a size approximately 10 % more than the size at which wild type cells divide. In cells longer than 15 µm, the rate of global translation reduced and then plateaued at lengths above about 18 µm (Figure 2H, I). In contrast, in an asynchronous population of *cdc25-22 hENT1 hsvTK* cells grown at a semi-permissive temperature of 30 °C, global transcription decreased only slightly with cell length up to lengths of 22 µm around 60 % longer than dividing wild type cells (Figure 2J, K). A decrease in transcription was reported in a population of enlarged fission yeast cells blocked in the cell cycle progression, but these cells were larger being over twice the size of dividing wild type cells [34]. Therefore, the global transcription machinery is not saturated in cells up to 22 µm wilst the rate of translation plateaus at 18 µm. We conclude that the plateau of global translation is unlikely to be due to transcription becoming limiting.

### Global cellular translation from G1 to G2

We next sought to determine how global translation and transcription scale with the increase in the amount of DNA at S-phase and with the cellular changes happening as cells proceed through mitosis and cell division. Wild type *S. pombe* cells spend the majority of their cell cycle in G2. The G1 phase is short so that DNA replication starts soon after completion of mitosis and is mostly completed by the time septated cells split to form two daughter cells [33]. This is also the case for the *hENT1 hsvTK* strain (Figure S3A). To assay protein and RNA synthesis in exponentially growing populations with cells of overlapping sizes in G1 and G2, and to assess cell-cycle effects in cells of the same size, we used the *cig1Δ cig2Δ puc1Δ* (*CCPΔ*) strain. This mutant strain has a delayed and more variable onset of S-phase compared to wild type cells [36] (Figure 3A, S3A). This means that the cell population has cells that are of the same size but which are located in the G1 or G2 phases of the cell cycle. *CCPΔ* cells were assayed for global translation and their DNA was stained with Nuclear-ID Blue. The maximum DNA concentration was determined within each cell and was used to classify cells as having either 1C, or 2C DNA content (Figure 3B). Cell lengths were also measured. In both the 1C and the 2C DNA content subpopulations, the median global translation per cell increased with cell size. The median global translation in cells of similar length increased by 35-40% in the G2 cells with a 2C DNA content compared with G1 cells with a 1C DNA content (Figure 3C, D).

**Figure 3.**
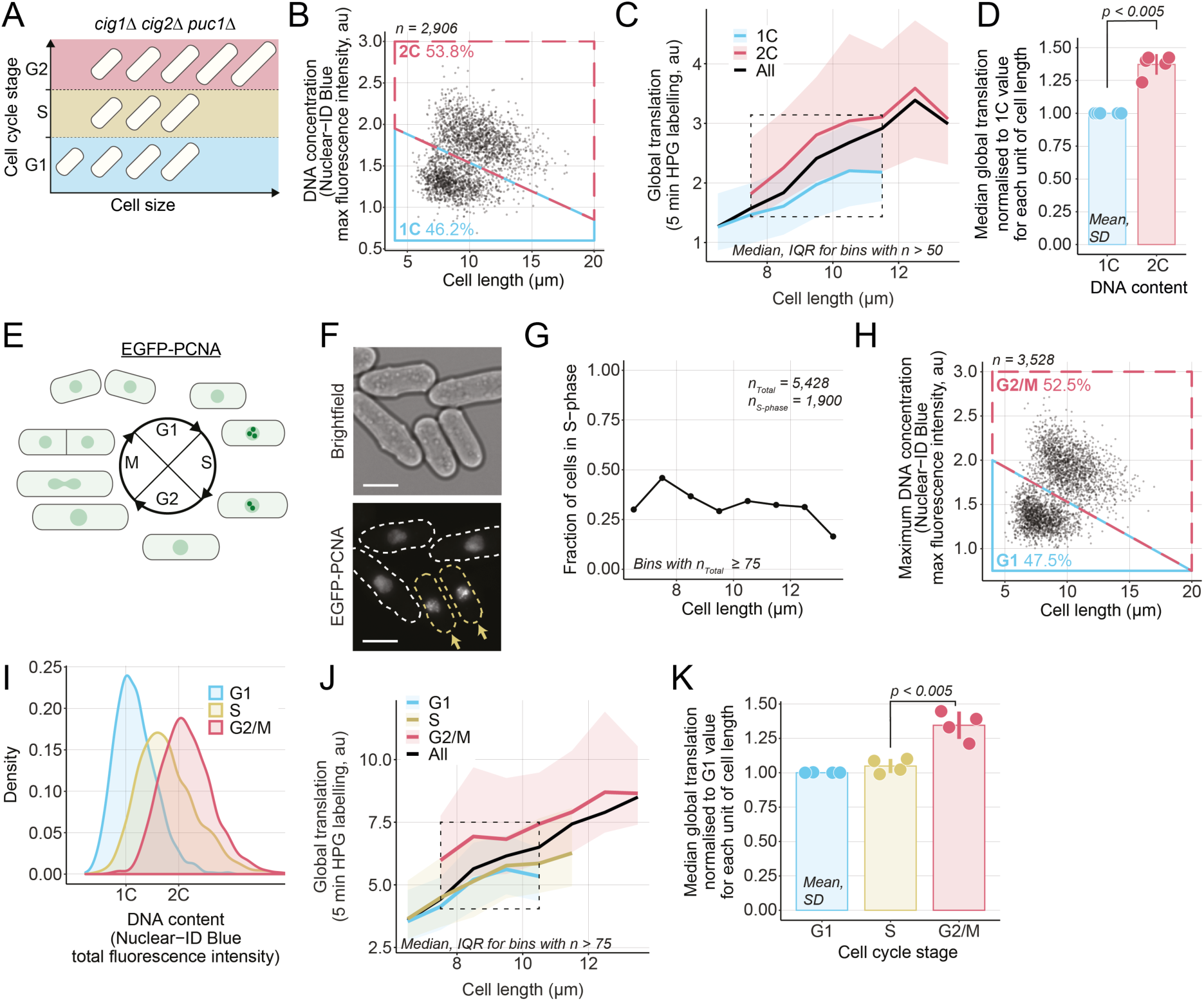
Global cellular translation from G1 to G2 in *CCPΔ* cells. (**A**) Schematic representation of the G1, S, and G2 subpopulation with overlapping cell sizes in the *CCPΔ* strain. (**B**) *CCPΔ* cells were assayed for global cellular translation. The maximum DNA concentration, measured as the maximum fluorescence intensity of the Nuclear-ID Blue stain in a cell, and cell length are used to categorise cells as having either 1C (blue box) or 2C DNA (red box). The percentage of cells in each category in shown. (**C**) Cells shown in (B) are grouped in length bins of 1 µm. Medians of global cellular translation (solid lines) and IQR (shaded areas) are shown for 1C (blue) and 2C DNA (red) subpopulations. The dashed line box marks the length bins which have both a 1C and a 2C median global cellular translation values. Bins containing more than 50 cells are shown. (**D**) For each of the 5 length bins boxed in (C), both 1C and 2C medians are normalised to their respective 1C global cellular translation values. Mean and SD of the normalised values (dots) are shown. For each DNA content, the normalised values (dots) are in the same order (left to right) as their corresponding length bins in (C). The *p*-value is calculated using a Welch’s unequal variances *t*-test. (**E**) Schematic of EGFP-PCNA localisation through the cell cycle. (**F**) Example images of brightfield and EGFP-PCNA fluorescence in *CCPΔ EGFP-pcn1* cells. The dashed lines in the EGFP-PCNA channel delimit the cell masks generated from the brightfield image. Cells with visible foci in the EGFP-PCNA channel are highlighted in yellow and marked with arrows. Scale bars represent 5 µm. (**G**) *CCPΔ* cells were assayed for global cellular translation using a 5-minute HPG incubation. Cells with EGFP-PCNA foci were identified by eye and binned in 1 µm intervals to compute the fraction of cells in S-phase per cell length. (**H**) The maximum DNA concentration and cell length were used to categorise cells not identified as in S-phase in (G) has either in G1 (blue box) or G2/M (red box). The percentage of cells in each category in shown. (**I**) Distribution of total DNA content of the cell populations categorised in (G) and (H) as measured per total Nuclear-ID Blue fluorescence intensity per cell. (**J**) Cells shown in (G) and (H) are grouped in length bins of 1 µm. Medians of global cellular translation (solid lines) and IQR (shaded areas) are shown for G1 (blue), S (yellow), and G2/M (red) subpopulations. The dashed line box marks the length bins which have G1, S, and G2/M median global cellular translation values. Bins containing more than 75 cells are shown. (**K**) For each of the 3 length bins boxed in (J), G1, S, and G2/M medians are normalised to their respective G1 global cellular translation value. Mean and SD of the normalised values (dots) are shown. For each cell cycle stage, the normalised values (dots) are in the same order (left to right) as their corresponding length bins in (J). The *p*-value is calculated using a Welch’s unequal variances *t*-test.

To understand further the increase in global translation from G1 to G2, we identified the S-phase subpopulation using a strain containing PCNA fused to an EGFP fluorescence marker [37]. During S-phase, EGFP-PCNA forms foci on replicating DNA so that cells in S-phase can be identified using fluorescence microscopy [38] (Figure 3E, F). The population of cells identified with EGFP-PCNA foci almost entirely overlapped with the population of cells replicating their DNA when assayed using 5-ethynyl-2’-deoxyuridine (EdU) (Figure S3B-D) indicating that the presence of EGFP-PCNA foci reliably identifies S-phase cells. These *CCPΔ EGFP-pcn1* cells were assayed for global translation. We identified cells with EGFP-PCNA foci (Figure 3G) and classified the remaining cells as in G1 or in G2/M based on their DNA concentration and cell length (Figure 3H). The distributions of total fluorescence intensity per cell of Nuclear-ID Blue are similar to the distributions of DNA content in the three populations indicating a reliable attribution of cell cycle stages (Figure 3I). Global translation was observed to increase with cell length in all subpopulations (Figure 3J). For a given cell length, global translation increased from the S to the G2/M subpopulation by about 30-35 %, but by less than 5 % from the G1 to the S-phase subpopulation (Figure 3J, K). This indicates that on transition of cells from S to G2 there is about a one third increase in the rate of translation.

### Global cellular transcription from G1 to G2

To understand how changes related to the G1, S, and G2 phases affect global transcription, we assayed *CCPΔ EGFP-pcn1 hENT1 hsvTK* cells for global transcription using a 20-minute EU incubation to compensate for their lower signal production. We identified cells with EGFP-PCNA foci (Figure 4A) and classified the remaining cells as in G1 or in G2/M based on their DNA concentration and cell length (Figure 4B). Global transcription increased for a given cell length from the G1 to the G2/M subpopulation by around 30-35 % and the S-phase subpopulation was found to have an intermediary global transcription value between the G1 and G2/M subpopulations of around 20-25 % (Figure 4J, K). This indicates that global transcription increases through S-phase, scaling approximately with the amount of DNA. An increase in global transcription with cell length and from G1 to G2 was also observed in a strain without the EGFP-PCNA marker (Figure S4A-C).

**Figure 4.**
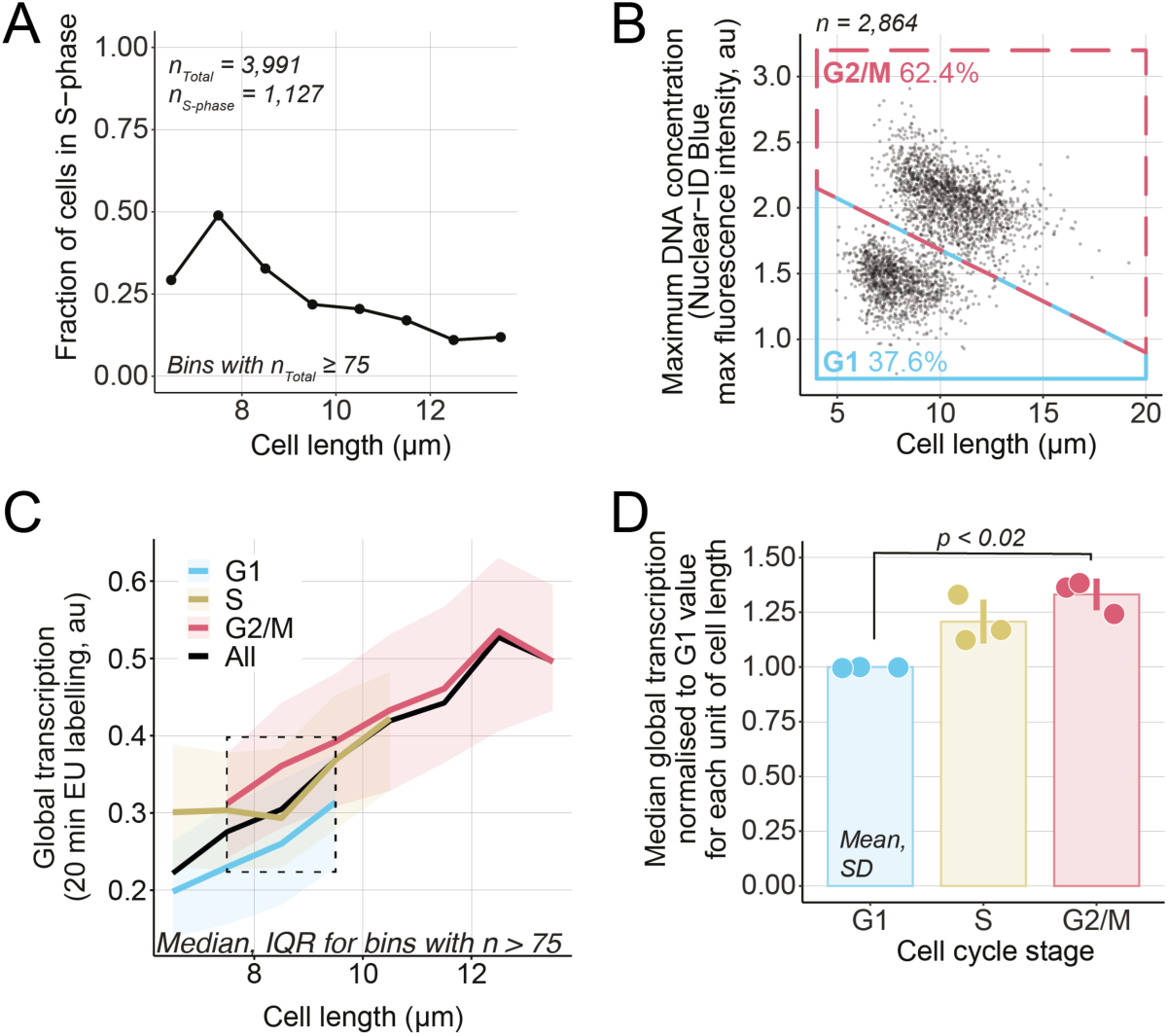
Global cellular transcription from G1 to G2. (**A**) *hENT1 hsvTK CCPΔ EGFP-pcn1* cells were assayed for global transcription using a 20-minute (almost linear) EU incubation. Cells with EGFP-PCNA foci were identified by eye and binned on 1 µm intervals to compute the fraction of cells in S-phase per cell length. (**B**) The maximum DNA concentration and cell length were used to categorise cells not identified as in S-phase in (A) as either in G1 (blue box) or G2/M (red box). The percentage of cells in each category in shown. (**C**) Cells shown in (A) and (B) are grouped in length bins of 1 µm. Medians of global cellular transcription (solid lines) and IQR (shaded areas) are shown for G1 (blue), S (yellow), and G2/M (red) subpopulations. The dashed line box marks the length bins which have G1, S, and G2/M median global transcription values. Bins containing more than 75 cells are shown. (**D**) For each of the 3 length bins boxed in (C), G1, S, and G2/M medians are normalised to their respective G1 global transcription value. Mean and SD of the normalised values (dots) are shown. For each cell cycle stage, the normalised values (dots) are in the same order (left to right) as their corresponding length bins in (C). The *p*-value is calculated using a Welch’s unequal variances *t*-test.

Thus, both global translation and global transcription increase from G1 to G2. Tanslation increases at the S/G2 transition or early in G2 and so is likely to be due to a subsequent cell cycle event dependent upon S-phase, whilst transcription increases throughout S-phase.

### Global cellular translation and transcription at mitosis

Next, we determined the dynamics of global translation and transcription at mitosis. To identify mitotic cells, we used strains expressing synCut3-mCherry, a truncated version of the condensin subunit Cut3 fused to the mCherry fluorescent reporter [39]. The synCut3-mCherry fusion protein is localised in the cytoplasm through interphase and rapidly accumulates in the nucleus at mitotic onset before being exported back to the cytoplasm from anaphase A onwards [39] (Figure 5A, B). Thus, in an asynchronous population, the progression through mitosis of each cell can be assessed based on the localisation of the synCut3-mCherry fluorescence signal and the number of nuclei in the cell. Uninucleated and binucleated cells with low nuclear synCut3-mCherry are in interphase, uninucleated cells with high levels of nuclear synCut3-mCherry are in mitosis between mitotic onset and anaphase A, and binucleated cells with high nuclear synCut3 are post-anaphase A. We assayed global translation in a *synCut3-mCherry* population and classified uninucleated and binucleated cells as having high or low nuclear synCut3 using their mean and median synCut3-mCherry fluorescence intensity (Figure 5C). We observed changes associated with the progression of cells into and through mitosis. For a similar cell size, global translation increased around 10 % early in mitosis, and decreased by about 20 % after anaphase A to below pre-mitotic levels (Figure 5D, E). Thus, global translation increases and then decreases as cells proceed through mitosis. In contrast, when we assayed global transcription in the *synCut3-mCherry hENT1 hsvTK* strain and categorised cells as uninucleated or binucleated and having either high or low nuclear synCut3-mCherry (Figure 5F), we found no change in global transcription for a given cell length in the different mitotic subpopulations (Figure 5G). This suggests that global transcription is not affected by the cellular changes happening in mitosis.

**Figure 5.**
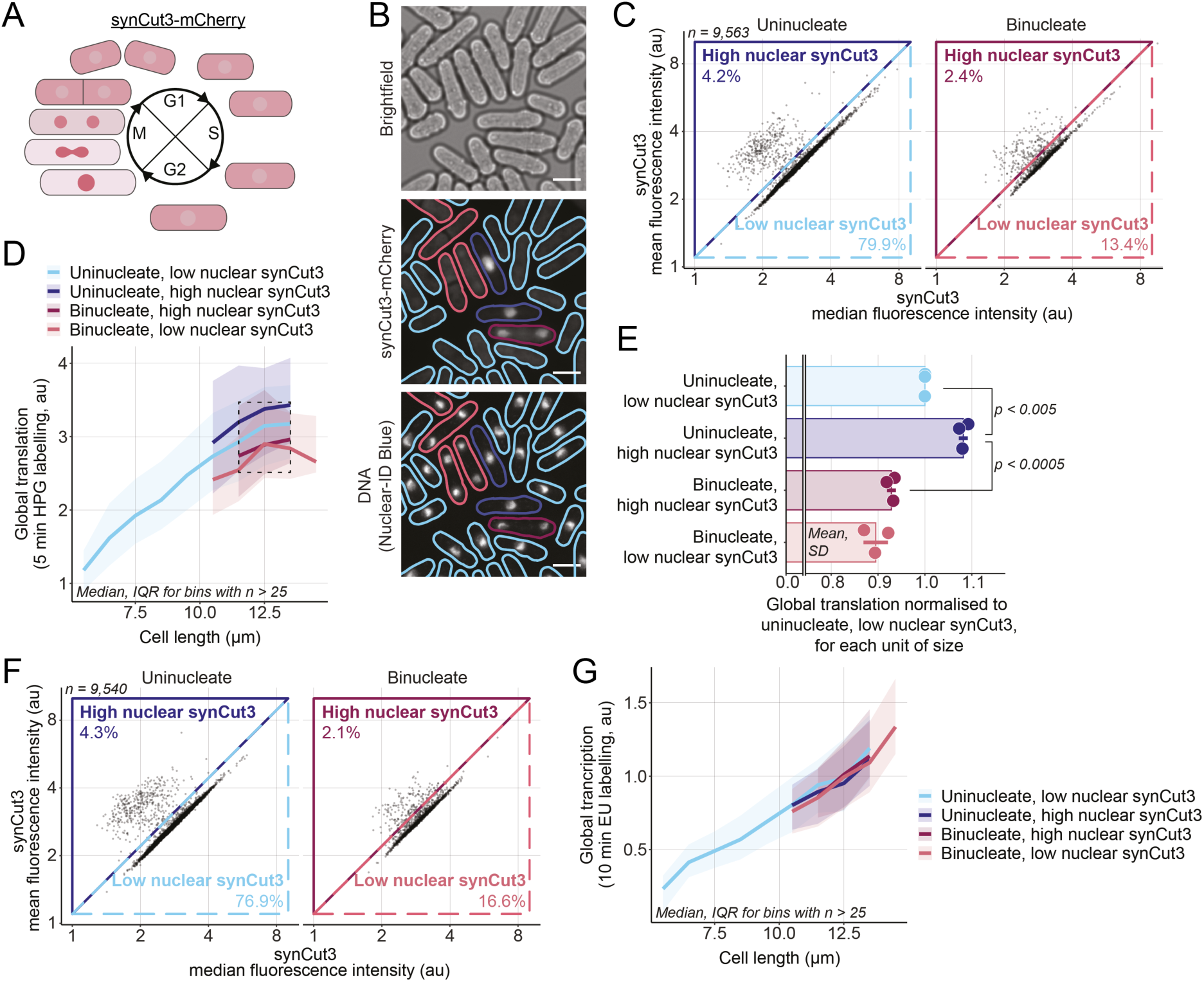
Global cellular translation and transcription at mitosis. (**A**) Schematic of synCut3-mCherry localisation through the cell cycle. (**B**) Example images of brightfield, synCut3-mCherry, DNA (Nuclear-ID Blue) fluorescence in *synCut3-mCherry* cells. The solid lines in the synCut3-mCherry and DNA channels delimit the cell masks generated from the brightfield image and is coloured according to the classification used in (D). The outlines of uninucleates are blue, binucleates are red, cells with high nuclear synCut3-mCherry have dark outlines, and cells with low nuclear synCut3-mCherry have light outlines. Scale bars represent 5 µm. (**C**) A population of *synCut3-mCherry* cells was assayed for global cellular translation. The mean and median whole-cell fluorescence synCut3-mCherry intensities of uninucleates and binucleates were used to categorise cells as having low/high nuclear synCut3-mCherry. The lines represent the delimitation of the different categories, the percentage of cells from the total population in each category is shown. (**D**) Cells shown in (C) are grouped in length bins of 1 µm. Medians of global cellular translation (solid lines) and IQR (shaded areas) are shown for uninucleates (blue) and binucleates (red), and with low (light) and high (dark) nuclear synCut3-mCherry signal subpopulations. The dashed line box marks the length bins which have median global cellular translation values for all 4 subpopulations. Bins containing more than 25 cells are shown. (**E**) For each of the 3 length bins boxed in (D), the medians of each subpopulation are normalised to their respective “uninucleate, low nuclear synCut3-mCherry” global cellular translation value. Mean and SD of the normalised values (dots) are shown. For each mitotic stage, the normalised values (dots) are in the same order (left to right) as their corresponding length bins in (D). The *p*-value is calculated using a Welch’s unequal variances *t*-test. For visual clarity, synCut3-mCherry is shortened to synCut3 in the figure. (**F**) Same as (C) for *synCut3-mCherry hENT1 hsvTK* cells assayed for global transcription. (**G**) Cells shown in (F) are grouped in length bins of 1 µm. Medians of global cellular transcription (solid lines) and IQR (shaded areas) are shown for uninucleates (blue) and binucleates (red), and with low (light) and high (dark) nuclear synCut3-mCherry signal subpopulations. Bins containing more than 25 cells are shown.

## Discussion

The rate of global transcription and to a lesser extent of translation have been investigated during the cell cycle of various eukaryotes [9,19–26,34,40], but the outcomes of these experimental investigations have been inconsistent with one another. This is probably due to effects of the different methods of synchronisation, perturbations due to a lack of steady state growth, and possibly variations between organisms and cell types. In this work, we use a single-cell approach to generate thousands of measurements of cell size, cell cycle stage, and global cellular translation and transcription, investigating unperturbed, steady-state, exponentially growing fission yeast cells.

The rate of global cellular translation increases linearly with cell size in wild type cells but plateaus at larger sizes. It is unclear what factor(s) may become limiting for global cellular translation in these larger cells. However, since global cellular transcription increases with cell size and but does not plateau within the range of sizes assayed, the plateau in global cellular translation is unlikely to be due to RNA becoming limiting. This is consistent with previous work suggesting that growth is mainly driven by the number of active ribosomes in cells [41, 42] and that cells enlarged beyond wild type sizes using cell cycle arrests experience cytoplasmic dilution of their proteins [4]. The increase in global cellular translation at the S/G2 transition and at the beginning of mitosis, and the decrease later in mitosis, suggest that there is regulation of global cellular translation as cells proceed through the cell cycle. This is consistent with recent work in synchronised mammalian cells showing an increase in translation early in mitosis followed by a decrease later [18]. Interestingly, proteins involved in translation initiation have been identified as substrates of the fission yeast cyclin dependant kinase (CDK1) Cdc2 [43] and CDK1 has been shown to phosphorylate the eukaryotic initiation factor 4E-binding protein (4E-BP1) in mammalian cell cultures [44].

The rate of global cellular transcription increases with cell size in both wild type cells, and in *cdc25-22* mutant cells which are up to 60 % larger, and the rate of transcription is increased in cells undergoing S-phase by 20 % and is 35 % higher in G2 cells which have completed S-phase, indicating that DNA content is limiting the global rate of transcription. Previous work has suggested that transcription by one of the RNA polymerases Pol II, increases with cell size [45–47]. This, in addition to the fact that global cellular transcription does not plateau in *cdc25-22* cells which are up to 60 % larger than wild type cells, suggests that Pol II and other RNA polymerases are not saturated in these enlarged cells. Therefore, the increase in global cellular transcription we observe from G1, through S-phase to G2, is unlikely to be the result of an increase in the amount of saturated DNA, but rather the result of an increase in the amount of unsaturated DNA leading to an increase in the probability of association of RNA polymerases with DNA. This is consistent with the dynamic equilibrium model for Pol II proposed for budding yeast [47]. The dynamic equilibrium model assumes that the increase in the occupancy of RNA polymerases is due to a dynamic equilibrium between free polymerases associating with the DNA and detaching from the DNA. We did not observe a reduction in global cellular transcription through mitosis unlike previous work in mammalian cells [48, 49] and budding yeast [50]. It is possible that undertaking mitosis without breaking down the nuclear envelope [51] prevents the reduction in transcription observed in mammalian cells undertaking an open mitosis. Another explanation might be that the larger genome of mammalian cells undergoes greater condensation for longer than is the case for fission yeast. The work carried in budding yeast was done using synchronised populations of cells so the difference observed might be the outcome of a perturbation as a consequence of synchronisation.

We propose that for the fission yeast, both translation and transcription steadily increase with cell size, but that the rate of translation becomes rapidly restricted when cells become larger than wild type dividing cells. This suggests that a component or components required for translation become limiting. It is unlikely that synthesis of RNA is the limiting factor since transcription still increases with size in cells larger than the wild type whilst translation does not. This may be related to the large resource and energy requirements of protein synthesis, meaning that there is only limited capacity for continued increase in the rate of translation. In cells dividing at wild type cell lengths, translation is regulated at different stages of the cell cycle; positively at the S/G2 transition and early in mitosis, and negatively later in mitosis. Perhaps changes in CDK activity through the cell cycle could influence the fraction of active ribosomes and be responsible for the cell cycle related changes in translation. Although the rate of transcription does not appear to be limiting in cells of this size, it is limited by DNA content. We suggest that global transcription is regulated by RNA polymerases which operate in dynamic equilibrium with DNA [47] and that global cellular translation is positively regulated in G2 possibly to coordinate with the increase in global cellular transcription that occurs during DNA replication.

Previous studies of global cellular translation and transcription during the cell cycle have given conflicting results. Our single cell approach gives us confidence that we have accurately described the changes of both translation and transcription with increasing cell size and progression through S-phase and mitosis/cell division in fission yeast cells. Knowledge of these changes is important for thinking about cellular control of macromolecular synthesis, and cell growth importance for the overall increase in cellular biomass. The approach we have used is employable with other eukaryotes to determine if there are conserved principles operating on these global cellular controls.

## Acknowledgements

We thank J. Curran, J. Greenwood, and E. Roberts for comments on the manuscript and N. Kapadia for sharing his plasmid. We also thank the Flow Cytometry and the Advanced Light Microscopy facilities at the Francis Crick Institute for their help with flow cytometry and microscopy, respectively. This work was supported by the Francis Crick Institute which receives its core funding from Cancer Research UK (FC01121), the UK Medical Research Council (FC01121), and the Wellcome Trust (FC01121). In addition, this work was supported by the Wellcome Trust Grant to PN [grant number 214183] The Lord Leonard and Lady Estelle Wolfson Foundation and Woosnam Foundation. For the purpose of Open Access, the author has applied a CC BY public copyright licence to any Author Accepted Manuscript version arising from this submission.

## Author contributions

Conceptualisation, C.B. and P.N.; Investigation, C.B.; Writing, C.B. and P.N.

## STAR Methods

### Key Resources table

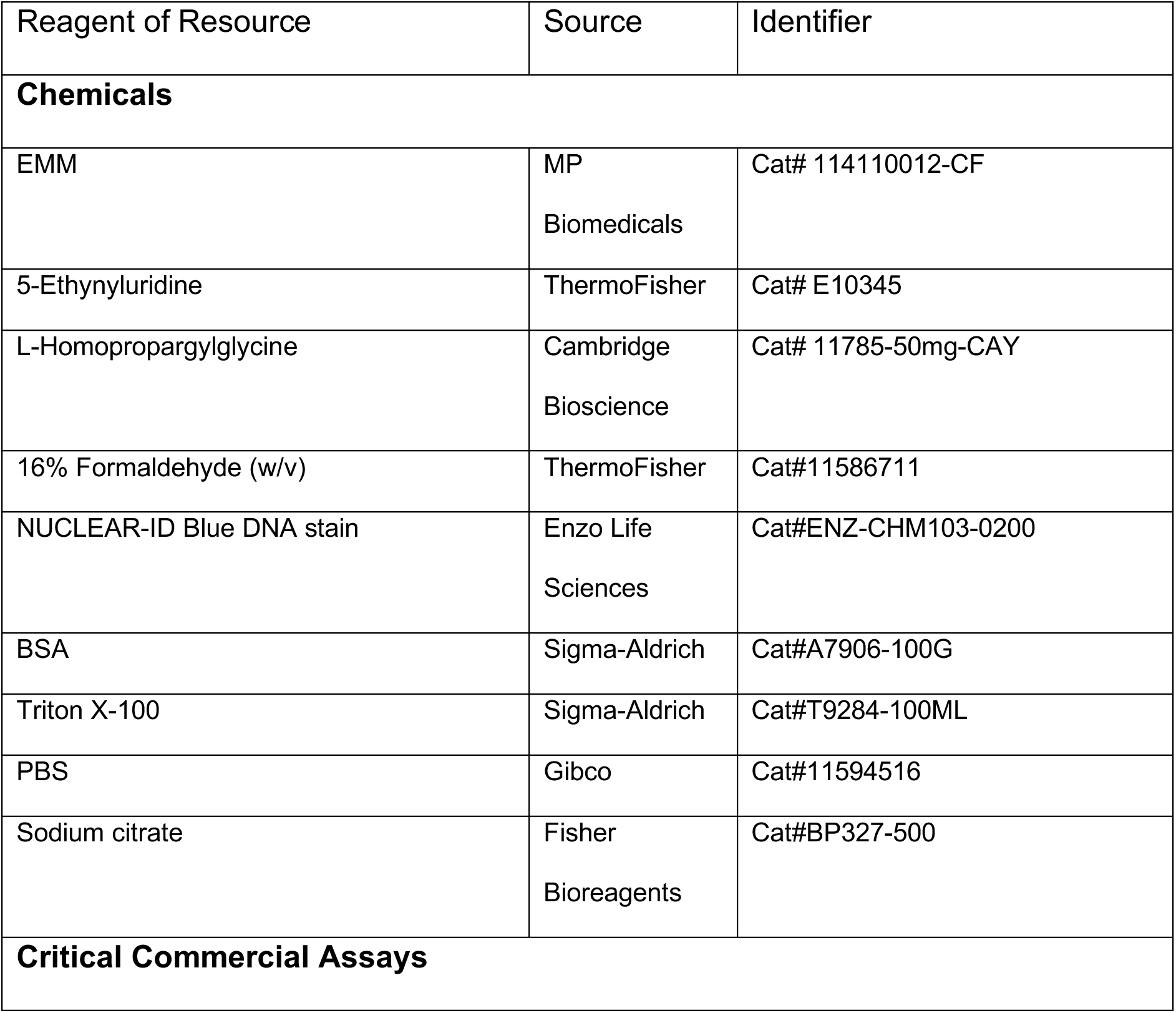
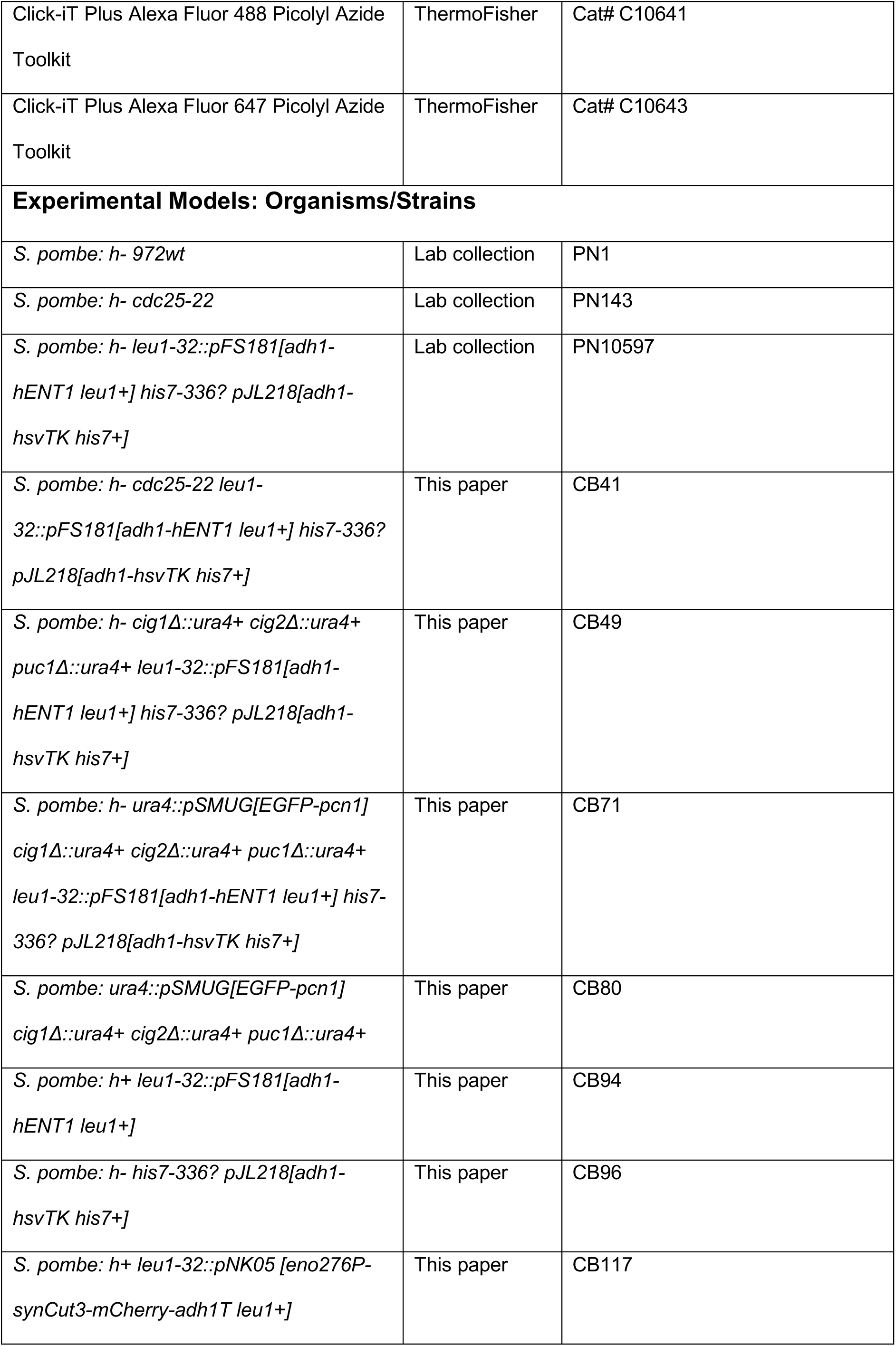
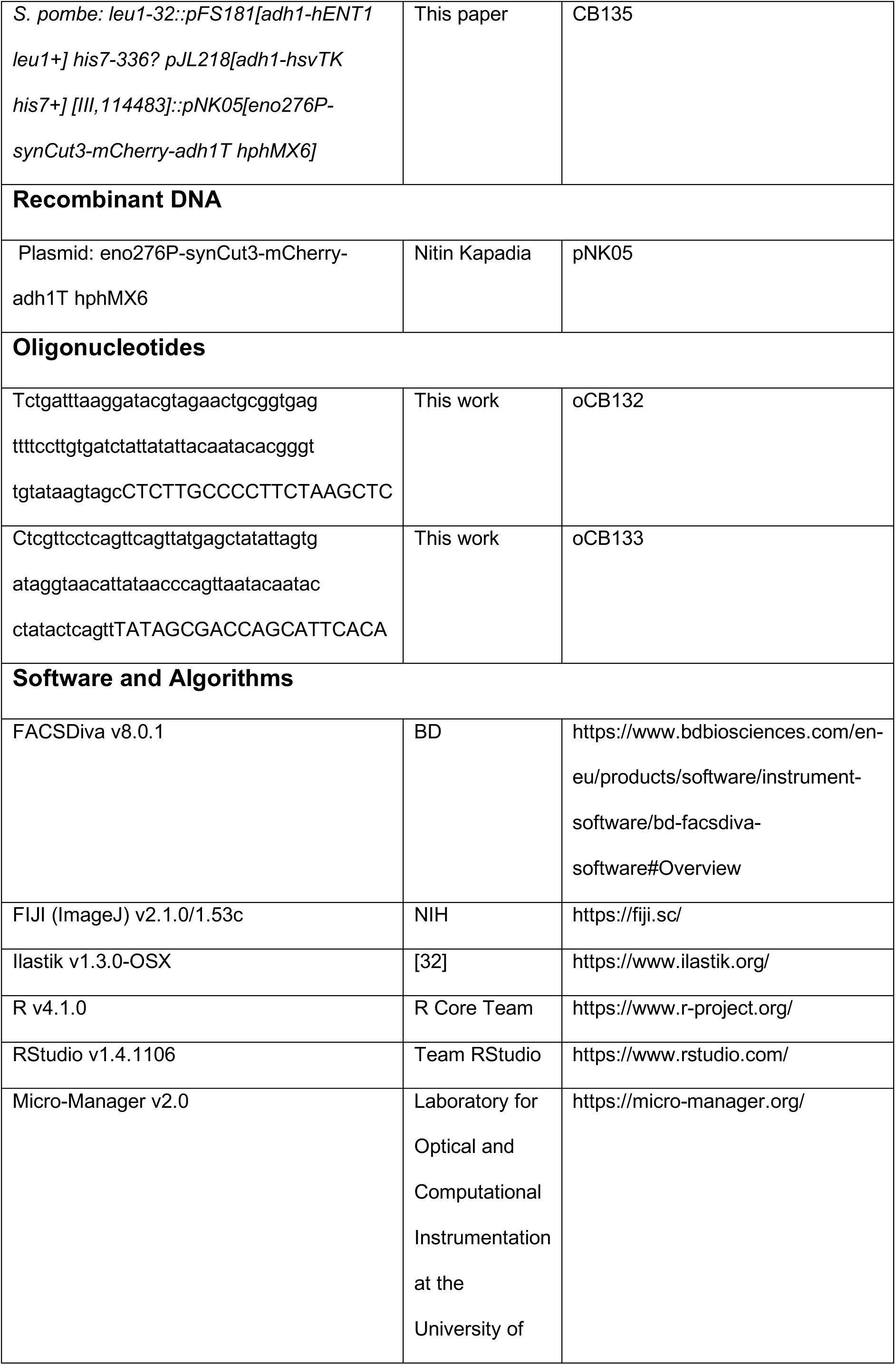
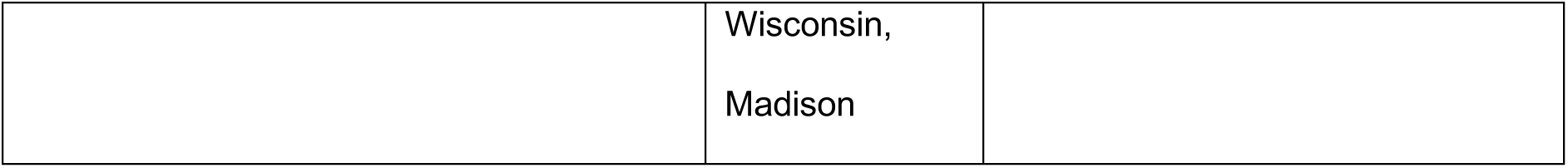

### Strain construction

All strains were constructed using random spore analysis after a genetic cross except for CB135 which was obtained by lithium acetate transformation of PN10597 with the [eno276P-synCut3-mCherry-adh1T hphMX6] construct from pNK05 amplified using the primers oCB132 and oCB133. All genotypes were confirmed phenotypically when possible (for temperature sensitive alleles and fluorescent markers) or by PCR for gene deletions.

### Cell cultures

Stationary cultures frozen and stored at −80 °C in 50% (v/v) YFM are patched on YE agar plates and incubated overnight (O/N) at 32 °C (or 25 °C if temperature sensitive). The patch is then streaked on a fresh EMM agar plate and cells are grown at 25 °C until visible single colonies form (typically around 4 days), a single colony is then patched on a fresh EMM agar plate and grown O/N at 25 °C. A 5 ml EMM liquid cultures is inoculated from a patch and grown in a stationary incubator O/N. The culture is then diluted in the morning in EMM to OD_595_ = 0.05 (calculated using an Amersham Ultraspec 2100 pro) in a flask and incubated for the day at 25 °C in a shaking incubator. The culture is then diluted in EMM to OD595 = 0.025 and grown O/N in a flask at 25 °C in a shaking incubator. In the morning, cells are diluted and used for the experiment.

### Global cellular transcription assay

An exponentially growing *S. pombe* culture of *hENT1 hsvTK* cells in EMM at 25 °C is diluted to 20 ml at OD595 = 0.3 (calculated using an Amersham Ultraspec 2100 pro) in a 50 ml flask, and placed in a shaking water bath for 1 h. Next, 4 µl of EU is added to the culture from a 100 mM stock solution in Milli-Q water to a final concentration of 20 µM. Immediately after addition of EU, a 3.84 ml sample of the culture is taken and fixed with 1.16 ml of a stock solution of 16 % (w/v) formaldehyde (methanol-free) in a 15 ml centrifuge tube, to a final concentration of 3.7 %, and vortexed for 5 s before being incubated at room temperature (19 to 23 °C) on a rocker, in the dark, for 40 min. This first sample will be used to compute the background signal. After 10 min, a second sample is taken from the culture and processed the same way, apart from being incubated for only 30 min. Fixed cells are then spun at 2,000 rcf for 5 min, and the supernatant is discarded. Cells are resuspended in 3 ml of PBS + 1 % (w/v) BSA, vortexed for 5 s, spun at 2,000 rcf for 5 min, and the supernatant is discarded. Cells are resuspended in in 6 ml of PBS + 1 % (w/v) BSA + 1 % (v/v) Triton X-100, vortexed for 5 s, and incubated at room temperature on a rocker for 30 min, in the dark. Cells are spun at 2,000 rcf for 5 min, the supernatant is discarded, and cells are resuspended in 6 ml of PBS + 1 % (w/v) BSA, vortexed for 5 s and incubated at room temperature on a rocker for 60 min, in the dark. Cells are spun at 2,000 rcf for 5 min, the supernatant is discarded, and cells are resuspended in 500 of 1X Click-iT reaction buffer (Thermo Click-iT Plus picolyl azide kit), and transferred to a 1.5 ml centrifuge tube. Cells are spun at 2,000 rcf for 5 min, the supernatant is discarded, and resuspended in 500 µl of the following reaction mix from the Thermo Click-iT Plus picolyl azide kit: 870 µl of 1X Click-iT reaction buffer (A), 10 µl of Alexa fluor at 500 µM (B), 15 µl of CuSO4 at 100 mM (C), 5 µl of Copper protectant (D), 10 µl of 10 X Click-iT buffer additive (E), 90 µl of Milli-Q water (F). To make the reaction mix, the solutions are added in the following order: A is mixed with B, C is mixed with D, E is mixed with F, AB is mixed with CD, EF is mixed with ABCD. Cells are incubated at room temperature on a shaker at 1,000 rpm for 30 min in the dark. Cells are then spun at 17,000 rcf for 15 s, the supernatant is discarded, and cells are resuspended in 800 µl of 50 mM sodium citrate and vortexed for 5 s. Cells are spun at 17,000 rcf for 15 s, the supernatant is discarded, cells are resuspended in 800 µl of 50 mM sodium citrate + 1:10,000 Nuclear-ID Blue, and vortexed for 5 s. Cells are spun at 17,000 rcf for 15 s, the supernatant is discarded, cells are resuspended in 800 µl of 50 µl sodium citrate. Cells are spun at 17,000 rcf for 1 s, the supernatant is discarded, cell are resuspended in 500 µl of 50 mM sodium citrate, and stored at 4 °C in the dark for 1 h before imaging.

### Global cellular translation assay

The protocol for the global translation assay is the same as the global transcription assay except the cells do not have the *hENT1* and *hsvTK* genes, and cells are incubated with 10 µM HPG for 5 min (4 µl from a 50 mM stock solution in Milli-Q water) instead of EU.

### Flow cytometry

Before running on the flow cytometer (BD LSRFortessa; excitation laser 488 nm, longpass filter 505 nm, bandpass filter 530/30 nm), samples are vortexed for 30 s, sonicated for 30 s (using a JSP Digital Ultrasonic Cleaner), and vortexed again for 30 s. The data is acquired using the BD FACSDiva (version 8.0.1) software. Single cells are gated based on their SSCA and FSCA profiles.

### Microscopy

All brightfield and fluorescence microscopy is performed using a Nikon Eclipse Ti2 inverted microscope equipped with Nikon Perfect Focus System, Okolab environmental chamber, and a Photometrics Prime Scientific CMOS camera. The microscope is controlled using the Micro-Manager v2.0 software. Fluorescence excitation is performed with a Lumencor Spectra X light engine fitted with the following excitation filters; 395/25 nm for imaging Nuclear-ID Blue; 470/24 nm for imaging EFGP, and Alexa Fluor 488; 575/25 nm for imaging mCherry; 640/30 nm for imaging Alexa Fluor 647. The emission filters used are the following: Semrock Brightline 438/24 nm for imaging Nuclear-ID Blue, Chroma ET525/50m for imaging EFGP, and Alexa Fluor 488; Semrock Brightline 641/75 nm for imaging mCherry; Semrock Brightline 680/42 nm for imaging Alexa Fluor 647. The dichroic mirrors used are the following: Semrock 409/493/573/652 nm BrightLine quad-edge standard epi-fluorescence dichroic beamsplitter for imaging Nuclear-ID Blue, EGFP, Alexa Fluor 488, Alexa Fluor 647; Chroma 59022bs dichroic beamsplitter for imaging mCherry. Images are taken using a Nikon Plan Apo 100x/1.45 Lambda oil immersion objective.

### Image segmentation and quantification

The brightfield images 1 µm below the focal plan of cells have a distinct outline and are therefore used to generate whole cell masks using Ilastik-1.3.0-OSX. The cell masks generated this way overlap well with the cells on the focal plane images. The 3 fluorescence images of the focal plane and the ± 0.5 µm z-stacks are maximum projected, all subsequent analysis is done on the maximum projected fluorescence images.

To generate the DNA masks, Ilastik-1.3.0-OSX is used on the Nuclear-ID Blue fluorescence images.

To obtain the number of nuclei per cell, the number of DNA masks within each whole cell mask is calculated using Fiji (ImageJ version 2.1.0/1.53c).

On all images, the scale is set using the function Analyze > Set Scale of Fiji so that the distance between 15.3609 pixels corresponds to 1 µm.

To generate single-cell measurements of cell length, the Analyze > Analyze particles function of Fiji is used on the whole cell masks to calculate for each mask; its Feret’s diameter (the longest distance between any two points within a mask, used as a measurement of cell length), its area, and its width (define as the width of the smallest rectangle enclosing the mask). Then the Analyze > Analyze particles function is used with the cell masks to calculate their corresponding fluorescence measurements on the fluorescence images, comprising of the total pixel intensity, mean pixel intensity, the median pixel intensity, and maximum pixel intensity. The masks are indexed so that the single-cell measurements of the different channels and the measurement of the number of nuclei are attributed to their corresponding cell mask.

The data is then processed using R (version 4.1.0) and RStudio (version 1.4.1106). For the global cellular transcription and translation signals, the median total fluorescence intensity of the background sample(s) within an experiment (cells immediately fixed after addition of EU, or HPG) is calculated. Then, the total fluorescence intensity of each cell is divided by the median background total fluorescence intensity. This allows all experiments to have fluorescence values roughly on the same scale and is convenient for processing.

Next, the background is subtracted based on cell length. Cells are grouped based on their length in bins spanning 1 µm (unless stated otherwise). For each length bin, the median background total fluorescence intensity is calculated on the background samples and subtracted to each cell total fluorescence intensity according to its length. The total fluorescence intensity of a cell normalised by the median background total fluorescence intensity, with the median total fluorescence intensity corresponding to its length then subtracted, is used as a measure of EU, or HPG incorporation

## Supplemental figure legends

**Figure S1.**
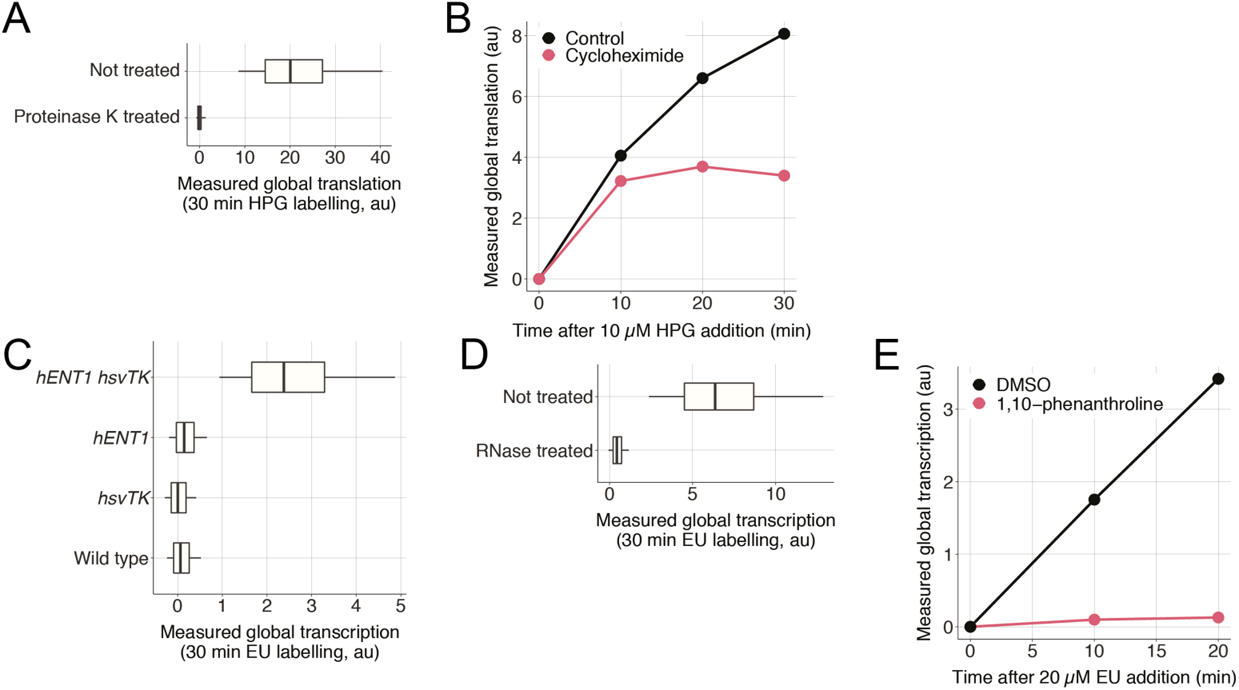
(**A**) Wild type cells were incubated for 30 minutes with 10 µM HPG, then assayed for global translation, then treated with 0.05 mg/ml Proteinase K at 55 °C for 4 h, and fluorescence was measured using flow cytometry. The 0.05, 0.25, 0.5, 0.75, and 0.95 population quantiles of at least 200,000 cells are shown. (**B**) Wild type cells were spun down and resuspended in EMM (control) or EMM + 10 mg/ml cycloheximide (t = 0), then assayed for global translation at different times using flow cytometry. Population medians of at least 200,000 cells are shown. (**C**) Cells expressing either *hENT1*, *hsvTK*, or both were pulsed with 10 µM EU for 30 minutes and assayed for global transcription using flow cytometry. The 0.05, 0.25, 0.5, 0.75, and 0.95 population quantiles of at least 200,000 cells are shown. (**D**) Cells expressing *hENT1* and *hsvTK* were pulsed with 10 µM EU and labelled with Alexa Fluor 488 azide, then treated with 0.1 mg/ml RNase A at 37 °C for 16 h, and the fluorescence signal was assessed using flow cytometry. The 0.05, 0.25, 0.5, 0.75, and 0.95 population quantiles of at least 200,000 cells are shown. (**E**) Cells expressing *hENT1* and *hsvTK* were pulsed with EU plus DMSO or EU plus 300 µg/ml 1,10-phenanthroline and global transcription was assayed at different time intervals using flow cytometry. Population medians of at least 200,000 cells are shown.

**Figure S2.**
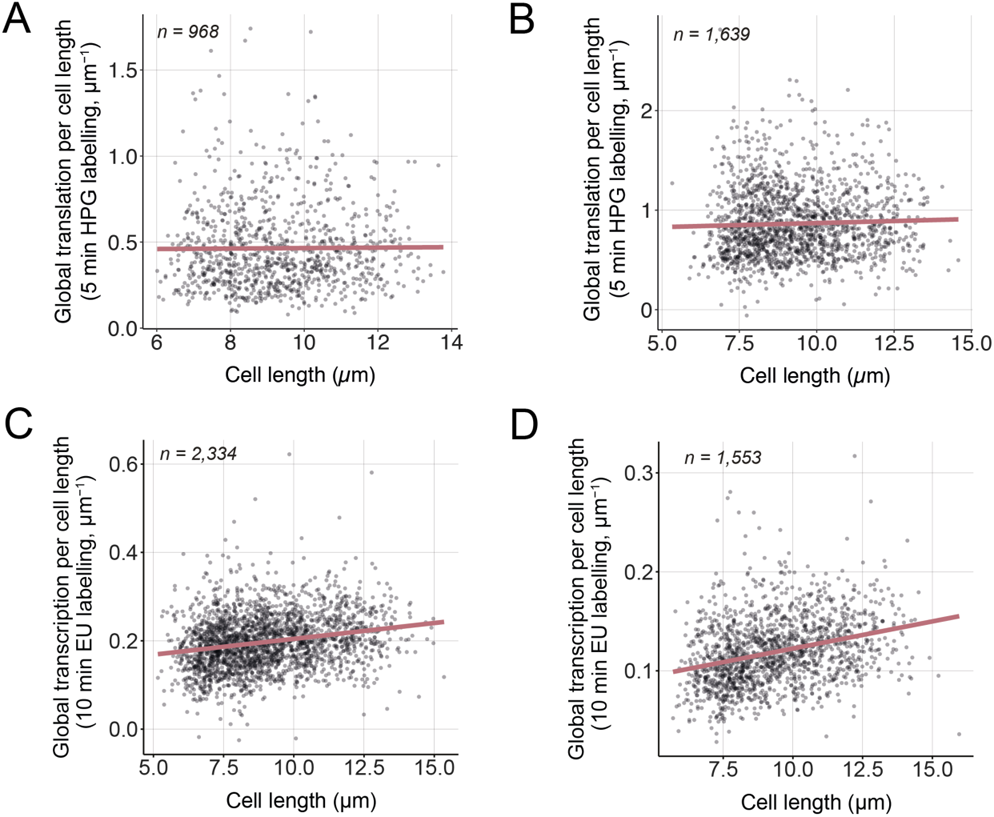
(**A**) and (**B**) Experimental replicates of Figure 2C. (**C**) and (**D**) Experimental replicates of Figure 2F.

**Figure S3.**
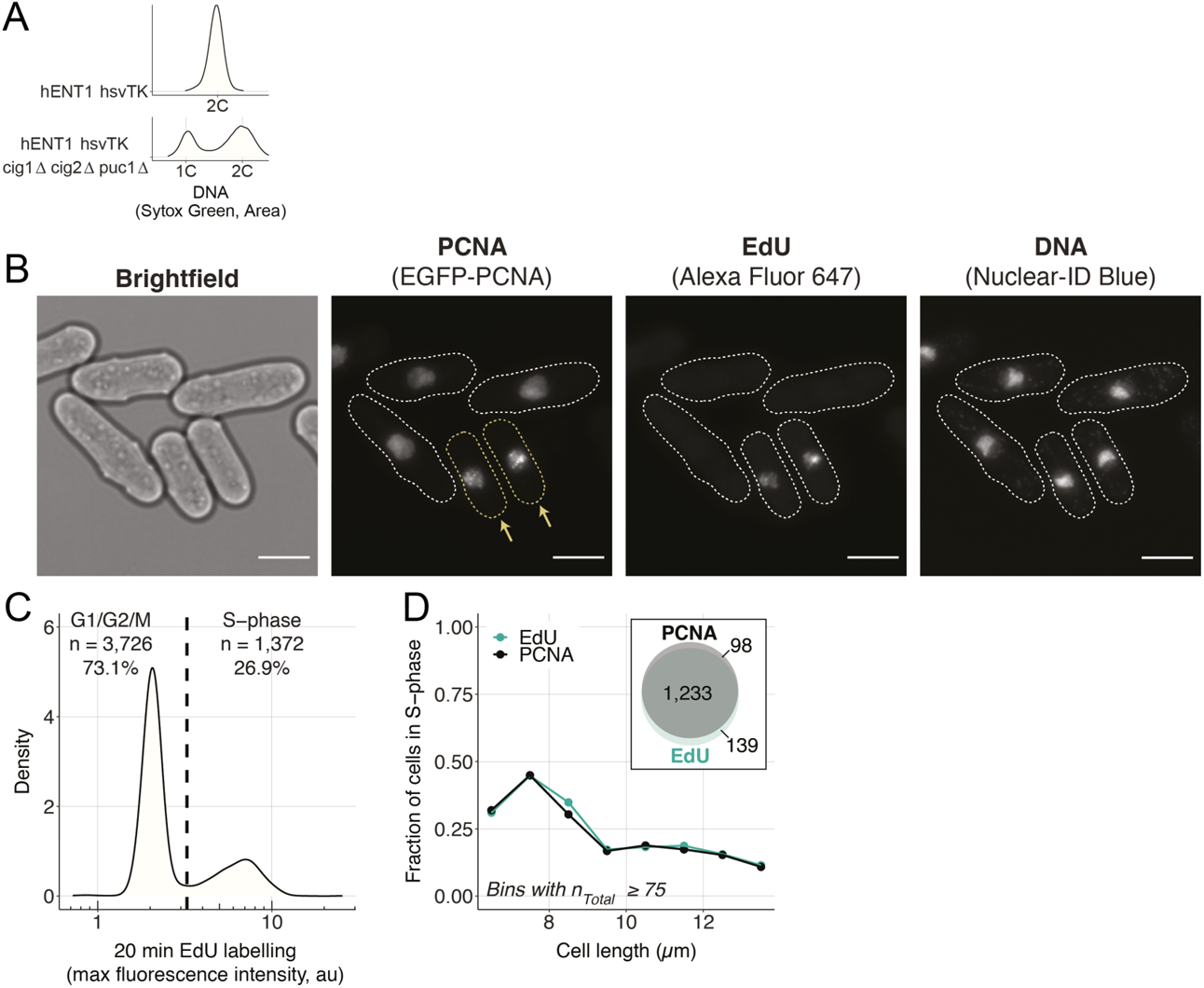
(**A**) Distribution of the amount of DNA in single cells in asynchronous populations, measured using the area of the fluorescence signal of Sytox Green by flow cytometry. For both populations, more than 200,000 cells were measured. Note that the 2C peak of the CCPΔ population is shifted to the right because the cells are longer and therefore have more mitochondrial DNA than the non-delete strain. (**B**) *hENT1 hsvTK EGFP-pcn1 CCPΔ cells* were pulsed with 200 µM EdU for 20 minutes and EdU incorporated in replicated DNA was fluorescently labelled using the same staining procedure used in the global transcription assay. Cells with visible foci in the PCNA channel are highlighted and marked with yellow arrows. The dotted white lines in the PCNA, EdU, and DNA channels delimit the cell masks generated from the brightfield image. The scale bar represents 5 µm. (**C**) Distribution of maximum fluorescence intensity of cells labelled with EdU. The dashed line represents the threshold (3.25 au) above which cells are considered in S-phase. (**D**) The fraction of cells in S-phase per cell length is computed using the EdU signal shown in (C), or the presence of EGFP-PCNA foci determined by eye. The inset shows the overlap in cell numbers between the two methods of identifying S-phase cells.

**Figure S4.**
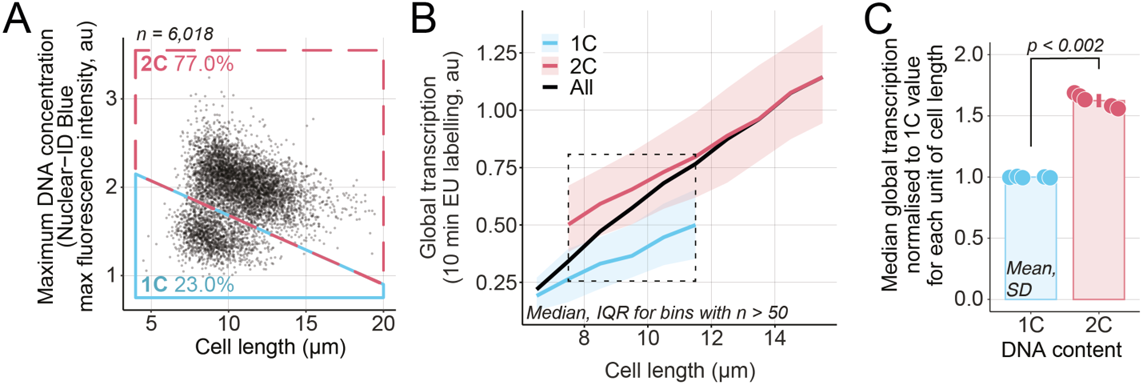
(**A**) *hENT1 hsvTK CCPΔ* cells were assayed for global transcription. The DNA concentration, measured as the maximum fluorescence intensity of the Nuclear-ID Blue stain in a cell, and cell length are used to categorise cells as having either 1C (blue box) or 2C DNA (red box). The percentage of cells in each box is shown. Black dots are single-cell measurements. (**B**) Cells are grouped in bins of 1 µm. Medians (solid lines) and inter quartile ranges (shaded areas) are shown for 1C (blue) or 2C DNA (red) populations. The dashed line box marks the length bins which have both a 1C and a 2C median global transcription values. (**C**) For each of the 5 length bins boxed in (B), both 1C and 2C medians are normalised to their respective global transcription 1C value. Mean and standard deviation of the normalised values are shown. The dots represent the median global transcription measurements per length bin boxed in (B). For each cell cycle stage, the normalised values (dots) are in the same order (left to right) as their corresponding length bins in (B). The p-value is calculated using a Welch’s unequal variances t-test.

